# Mapping the Host-Pathogen Space to Link Longitudinal and Cross-sectional Biomarker Data: *Leptospira* Infection in California Sea Lions (*Zalophus californianus*) as a Case Study

**DOI:** 10.1101/819532

**Authors:** K.C. Prager, Michael G. Buhnerkempe, Denise J. Greig, Anthony J. Orr, Eric D. Jensen, Forrest Gomez, Renee L. Galloway, Qingzhong Wu, Frances M.D. Gulland, James O. Lloyd-Smith

## Abstract

Confronted with the challenge of understanding population-level processes, disease ecologists and epidemiologists often simplify quantitative data into distinct physiological states (e.g. susceptible, exposed, infected, recovered). However, data defining these states often fall along a spectrum rather than into clear categories. Hence, the host-pathogen relationship is more accurately defined using quantitative data, often integrating multiple diagnostic measures, just as clinicians do to assess their patients. We use quantitative data on a bacterial infection (*Leptospira interrogans*) in California sea lions (*Zalophus californianus*) to improve both our individual-level and population-level understanding of this host-pathogen system. We create a “host-pathogen space” by mapping multiple biomarkers of infection (e.g. serum antibodies, pathogen DNA) and disease state (e.g. serum chemistry values) from 13 longitudinally sampled, severely ill individuals to visualize and characterize changes in these values through time. We describe a clear, unidirectional trajectory of disease and recovery within this host-pathogen space. Remarkably, this trajectory also captures the broad patterns in larger cross-sectional datasets of 1456 wild sea lions in all states of health. This mapping framework enables us to determine an individual’s location in their time-course since initial infection, and to visualize the full range of clinical states and antibody responses induced by pathogen exposure, including severe acute disease, chronic subclinical infection, and recovery. We identify predictive relationships between biomarkers and outcomes such as survival and pathogen shedding, and in certain cases we can impute values for missing data, thus increasing the size of the useable dataset. Mapping the host-pathogen space and using quantitative biomarker data provides more nuanced approaches for understanding and modeling disease dynamics in a system, yielding benefits for the clinician who needs to triage patients and prevent transmission, and for the disease ecologist or epidemiologist wishing to develop appropriate risk management strategies and assess health impacts on a population scale.

**Author Summary:** A pathogen can cause a range of disease severity across different host individuals, and these presentations change over the time-course from infection to recovery. These facts complicate the work of epidemiologists and disease ecologists seeking to understand the factors governing disease spread, often working with cross-sectional data. Recognizing these facts also highlights the shortcomings of classical approaches to modeling infectious disease, which typically rely on discrete and well-defined disease states. Here we show that by analyzing multiple biomarkers of health and infection simultaneously, treating these values as quantitative rather than binary indicators, and including a modest amount of longitudinal sampling of hosts, we can create a map of the host-pathogen interaction that shows the full spectrum of disease presentations and opens doors for new insights and predictions. By accounting for individual variation and capturing changes through time since infection, this mapping framework enables more robust interpretation of cross-sectional data; e.g., to detect predictive relationships between biomarkers and key outcomes such as survival, or to assess whether observed disease is associated with the pathogen of interest. This approach can help epidemiologists, ecologists and clinicians to better study and manage the many infectious diseases that exhibit complex relationships with their hosts.

## Introduction

To gain insights into population-level trends, disease biomarker data are often reduced to binary form (e.g. presence/absence of a pathogen, antibodies or disease) for statistical analyses and parameterizing models of disease transmission. By contrast, to understand disease in an individual, the full quantitative range of available biomarker information is used to determine the precise clinical status of an individual, make treatment decisions, assess prognoses and limit transmission risk to others. While clinicians consider a clinically ill individual with a high or rising antibody titer as diagnostic of a current or recent infection [1], ecologists or epidemiologists typically classify individuals as exposed or not, and infected or not, based on a cut-off titer value [2], potentially discarding useful information contained in finer scale variations in titer magnitude. More detailed data on infection status and health can provide key information to both the clinician and ecologist that can help with accurate diagnosis (clinician) and effective system conceptualization, model construction and parameterization (ecologist). However, such data can be difficult to interpret, particularly for wildlife hosts, and all data types may not be available for each individual assessed. Hence, cases captured in clinical and surveillance data often do not fit neatly into distinct categories. Severely ill, recently infected individuals are easily identified (e.g. by high antibody titer, pathognomonic clinical signs, detection of pathogen), but are often just the tip of the iceberg. In reality, a variety of presentations may exist at each point along the timeline from infection to recovery, with individuals exhibiting a range of disease severity and antibody titers (e.g. from severely ill to apparently healthy and with very high to undetectable titers), and with both infected and uninfected individuals detected at any given combination of disease severity and antibody titer (Fig 1 and S1 Box).

**Fig 1.**
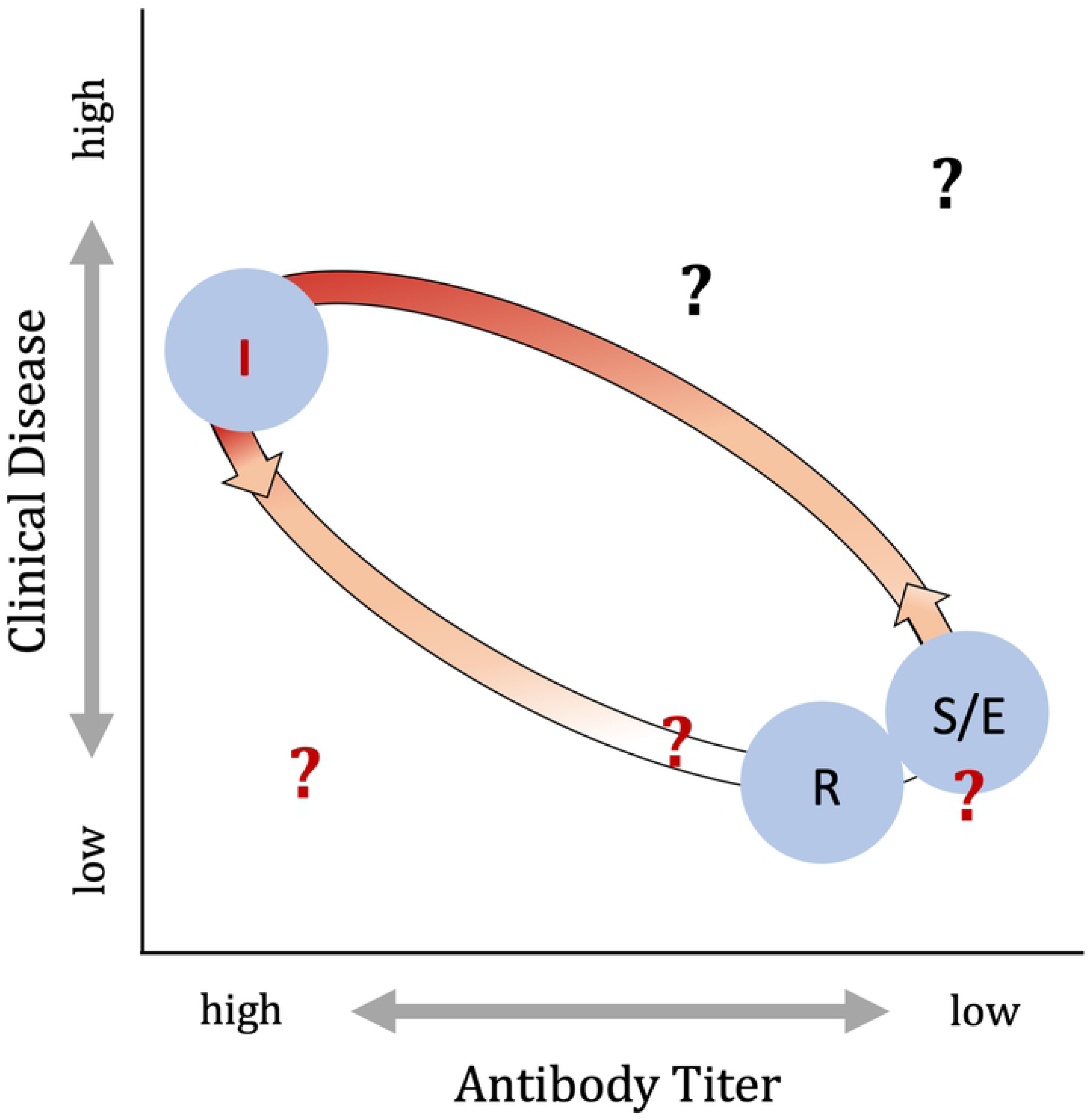
Map showing how infected individuals move through the “host-pathogen space” in dimensions reflecting severity of clinical disease (y-axis) and time since infection (∼antibody titer, x-axis). When susceptible individuals become infected (indicated by red shading – the intensity of shading indicates the probability that an assay for current infection – e.g. PCR – would be positive), they move through this space in a trajectory towards higher titer and more severe clinical disease states and then, depending on the type of pathogen and the host-pathogen interaction, they eventually return to a state of good health and their titers decline. The shape of this trajectory and their infection state will differ based on the host-pathogen system and the degree of heterogeneity in host responses. Here we show the trajectory that would be expected based on the canonical susceptible (S), exposed (E), infected/infectious (I) and recovered (R) model of infectious disease dynamics with the position of individuals as they would pass through each of the four states indicated by the blue circles. However, individuals – represented by question marks in this figure – experiencing more complex host-pathogen interactions may fall outside of this canonical trajectory (see S1 Box for further detail).

Recently, efforts have been made to assess how biomarkers of disease and infection change relative to each other and over time, with the aim of identifying consistent patterns to improve our understanding of host-pathogen dynamics in human [3] [4], domestic animal [5], experimental [3], and wildlife systems [6] [7]. Longitudinal studies in which individuals are monitored through time provide key insights into how specific host-pathogen biomarkers, e.g. antibody titer, measures of disease severity, and pathogen load, change through the course of infection and recovery [7–10], with some studies showing how biomarker values may be associated with specific outcomes such as survival and transmission [3, 11]. In systems for which biomarkers show predictable temporal variation, quantitative data may provide information about an individual’s stage in the infection and recovery process [3, 6, 12, 13], enhancing our understanding of population-level dynamics by providing key data for model structure and parameterization [6, 10, 12-19]. Assessment of quantitative values and multiple biomarkers can also elucidate individual within-host dynamics such as the outcome of an infection (infection chronicity, survival), how heterogeneity in antibody titer responses relates to clinical disease or symptoms, and probability of transmission to others [3, 11–13, 15, 20–22]. These findings can have direct implications on both the individual scale (e.g. triaging and treating patients, assessing prognosis and forward transmission risk) and the population scale (e.g. controlling transmission and hence outbreaks, predicting population dynamics, estimating incidence). These previous studies highlight the usefulness of including multiple data types, of understanding the nature of the relationship between multiple biomarkers of infection and disease, and of using quantitative data to better understand host-pathogen dynamics to make informed management decisions. However, although these studies explore facets of this new frontier in infectious disease dynamics, none combine all facets within a single study system, and few focus on disease in wildlife species.

We address this gap by linking longitudinal and cross-sectional data on multiple disease measures from an unconventional study system: *Leptospira interrogans* serovar Pomona (henceforth *“Leptospira*”) infection in California sea lions (*Zalophus californianus*). This system exhibits yearly, seasonal *Leptospira* outbreaks of varying magnitude, as reflected in both clinical cases of *Leptospira* infection seen at marine mammal rescue and rehabilitation centers [23] and in population-level serosurveys [24]. *Leptospira* is a good model for examining complex manifestations of a host-pathogen relationship, as mammals infected by pathogenic species within the genus *Leptospira* can exhibit a wide range of clinical presentations, from fulminant clinical disease to silent infections, and while some hosts may clear the infection quickly, others continue to shed the pathogen for months to years. The dominant clinical signs of leptospirosis (the disease caused by *Leptospira* infection) in California sea lions reflect the kidney damage inflicted by the bacteria, and clinically ill sea lions present in varying stages of renal failure. The host-pathogen relationship for pathogenic *Leptospira* spp. is conventionally attributed to specific *Leptospira* strain-host species pairs and described dichotomously, as an acute and potentially fatal infection in ‘accidental’ host species, or as a chronic and predominantly subclinical infection in ‘maintenance’ host species [1, 25]. Yet, California sea lions show characteristics of both accidental and maintenance hosts. During major outbreaks, roughly two-thirds of sea lions stranding with clinical *Leptospira* infections die – typical of accidental hosts. However, genetic evidence [26] and age-structured sero-epidemiology [24] suggest that *Leptospira* is enzootic in the sea lion population, and furthermore subclinical chronic infections – typical of maintenance hosts – occur in sea lions and are the possible mechanism for population-level pathogen persistence from one outbreak to another [19, 27, 28].

Using longitudinal data on antibody titer, disease severity and pathogen shedding, we track the temporal progression of *Leptospira* infections in California sea lions that experienced either severe illness or subclinical infection. We use the relationship between these different biomarkers to create a ‘host-pathogen space’ in which we track the progression of known infected individuals through time and establish that they follow a clear, unidirectional trajectory. Using this mapping approach, we then plot cross-sectional data from a broader group of sea lions – either apparently healthy, wild-caught individuals, or those stranding due to a broad range of health issues (i.e. not pre-selected for or against leptospirosis), and use the patterns cast by the longitudinal data to interpret those in the cross-sectional data. In human terms, the longitudinal data are akin to disease-specific long-term monitoring of individual cases, whereas the cross-sectional data are akin to prospective, random population surveillance, and unfiltered sampling of hospital patients, and are therefore more representative of the overall population. We show that the longitudinal data broadly capture the patterns in the cross-sectional data, suggesting consistency in dynamics despite the greater set of individual presentations present in the cross-sectional data. Our identification of a consistent trajectory through host-pathogen space enables us to roughly situate cross-sectionally sampled individuals in their time-course of infection, showing how our approach could elucidate disease dynamics in many systems – from wildlife to humans – where most available data are cross-sectional. We also find that patterns within the host-pathogen space provide population-level insights into the range of disease experienced, duration of shedding, and associations between antibody titer and infection status. This allows us to explore predictive relationships such as links between disease severity and survival, and between antibody titer and shedding duration. We also identify important differences between patterns in cross-sectional and longitudinal data, and generate and test hypotheses regarding the source of these differences, e.g., we identify renal disease from causes other than *Leptospira*.

## Results

### Establishing a Host-Pathogen Trajectory with Longitudinal Data

We tracked the temporal progression of three important biomarkers of *Leptospira* infection – anti-*Leptospira* serum antibody titer, renal compromise, and urinary leptospiral DNA shedding – in 15 sea lions that were followed longitudinally from infection to clinical recovery. Thirteen of these were initially severely ill and were followed for 6-12 weeks (henceforth termed CLINICAL), and 2 never showed clinical signs and were followed for 3 years (termed SUBCLINICAL for subclinical, or SUB1 and SUB2 when referred to individually; Table 1). Combined, data from the CLINICAL and SUBCLINICAL animals enabled us to assess host-pathogen dynamics in animals exhibiting a range of initial clinical disease. The CLINICAL animals are typical of what would be reported by hospitals or rehabilitation centers for a given disease but may comprise only a small fraction of infections experienced in a population. The majority of acute infections may involve no evident disease, similar to the SUBCLINICAL animals, and would only be detected through prospective surveillance efforts and unfiltered sampling of hospital cases.

**Table 1.** Description of the different data sets used in our study. Columns include the category of data “Group” and Sub-group”, the “Sample Size” of unique individuals, the “Selection Criteria” used for inclusion, the “Additional Details” regarding individuals included, the “Day 0”, i.e. the first day for which *Leptospira* infection related biomarkers were tracked in an individual, longitudinally monitored sea lion, and the “Length of Observation”, i.e. the period of time over which data were collected.

| Group | Sub-group | Sample Size | Selection Criteria | Additional Details | Day 0 | Length of Observation |
| --- | --- | --- | --- | --- | --- | --- |
| Longitudinal | CLINICAL | 13 | Presented initially with clinical signs of severe renal compromise consistent with leptospirosis*. | Survived infection and released into wild 6-12 weeks after admission to rehabilitation center. | First day anti-*Leptospira* antibody titer detected (0 – 18 days of admission) | 6 - 12 weeks |
| | SUB | 2 | Never showed clinical signs of leptospirosis. Admitted to rehabilitation center for treatment of other condition. Magnitude of the first detected anti-*Leptospira* antibody titers, and timing (October of a major *Leptospira* outbreak year in the wild sea lion population {Greig, 2005 #52}, suggest relatively recent *Leptospira* infection. | **SUB1:** No detectable anti-*Leptospira* antibodies initially, but seroconversion occurred (i.e. acquired anti-*Leptospira* antibodies) at some unknown point during rehabilitation, and in the absence of any observed clinical signs of leptospirosis. | First day anti-*Leptospira* antibody titer detected ($\log_2$ titer=10 on 10/23/11, 15 months after admission). | 3 years |
| | | | | **SUB2:** Moderately high anti-*Leptospira* antibody titer at admission. No clinical signs of leptospirosis. Released into wild 3 weeks after admission. Readmitted 3 months after initial admission, still no clinical signs of leptospirosis. | First day anti-*Leptospira* antibody titer detected ($\log_2$ titer=7 on 10/18/11, the day of admission) | |
| Cross-sectional | STRAND | 724 | All sea lions admitted to rehabilitation center for any cause, including leptospirosis. (i.e. not filtered by clinical signs) |  | N/A | 1 day |
|  | WILD | 730 | Apparently healthy, free-ranging sea lions. |  | N/A | 1 day |

\* Leptospirosis is the disease caused by infection with pathogenic species within the genus *Leptospira*.

We tracked changes in clinical disease using a ‘renal index’ that we derived from serum chemistry values (i.e. blood urea nitrogen, creatinine, sodium, chloride and phosphorus) associated with the compromised renal function seen in severe cases of leptospirosis [23]. Within the first 72 hours of admission to rehabilitation the severely ill animals that survived (CLINICAL) had high initial renal index values that ranged from 4.15 to 13.67, but they recovered rapidly with all scores declining into the healthy range within 15 to 61 days (median = 27 days; Fig 2A). By contrast, in the three years that they were monitored, we never detected serum chemistry evidence of renal compromise in the subclinical animals (SUB1 and SUB2; Fig 2B).

**Fig 2.**
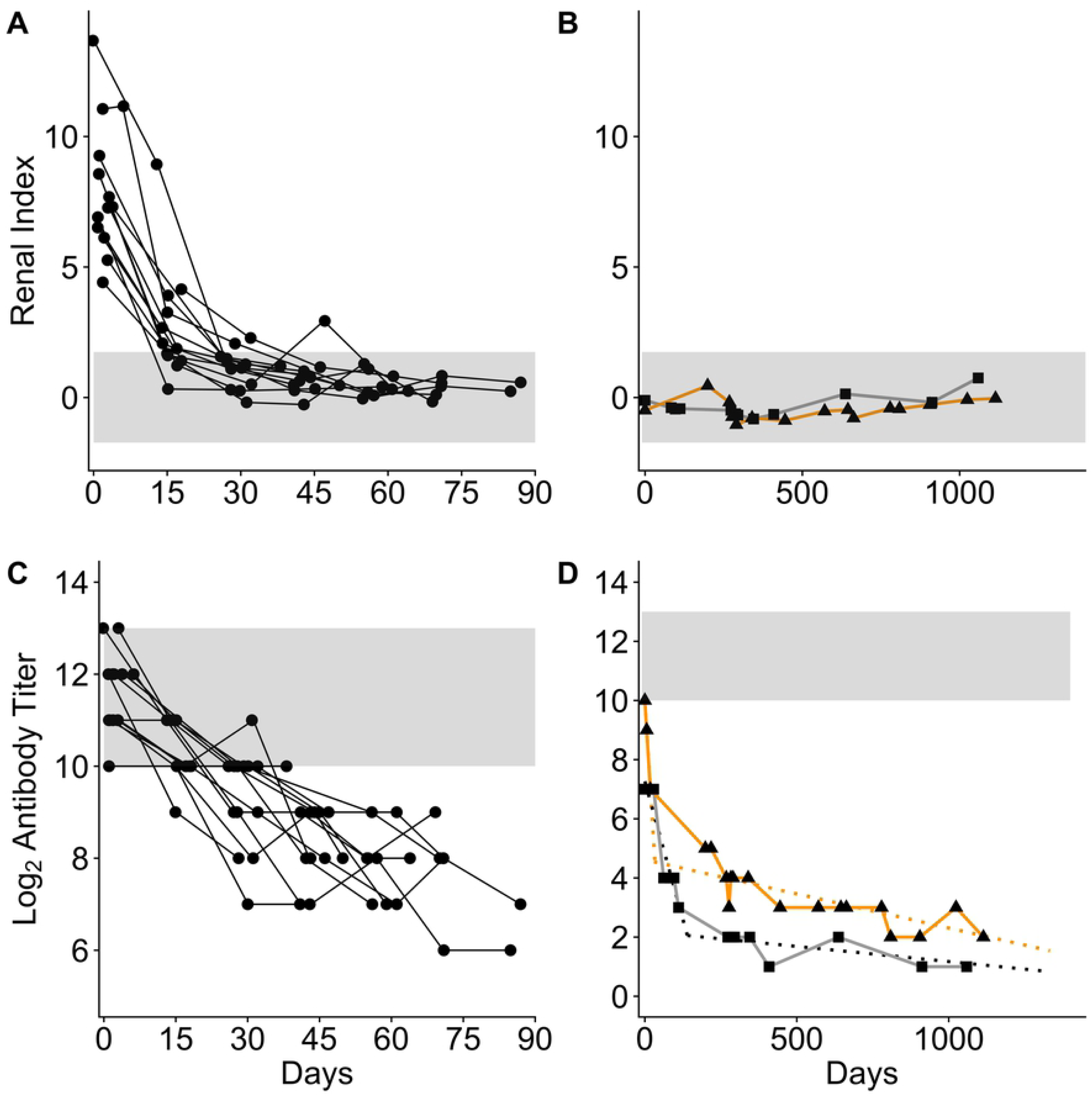
Changes in antibody titer and renal index in longitudinally sampled California sea lions. Renal index scores (A) and log_2_ anti-*Leptospira* antibody titer (C) by time in individual sea lions that stranded with clinical signs of leptospirosis and were followed for 6 – 12 weeks (CLINICAL dataset). Renal index scores (B) and log_2_ antibody titer (D) by time in two stranded sea lions – SUB1 (square, grey line) and SUB2 (triangle, orange line) – that never showed *Leptospira-*related clinical disease and were monitored for 3 years (SUB dataset). In panel D, regression lines, as determined by piecewise linear regression, are drawn through first and second phases of antibody titer decline for each SUB animal. For CLINICAL animals, day 0 is the day of admission to rehabilitation, for SUB1 and SUB2, day 0 is the day when anti-*Leptospira* antibodies were first detected. Grey horizontal bands in panels C and D delineate the full range of initial antibody titers in the CLINICAL animals, and in panels A and B they delineate the 95% interquantile range of renal index scores in apparently healthy, uninfected, seronegative wild-caught animals.

Antibody titers in individual CLINICAL sea lions exhibited simple exponential decay (Fig 2C), while the SUBCLINICAL animals exhibited a more complex pattern. Visual inspection of the SUBCLINICAL data suggested a biphasic pattern with an initial rapid phase consistent with that of the CLINICAL animals, followed by much slower decay (Fig 2D). Using a simple linear regression for each individual, we calculated half-life (t_1/2_) estimates in CLINICAL sea lions that ranged from 6.4 to 29.4 days with a median t_1/2_ of 17.1 days (Table 2). Using piecewise linear regression we calculated first phase t_1/2_ values of 26.8 and 6.1 days for SUB1 and SUB2 respectively, and second phase values of 976 and 433 days (Table 2). The fact that first phase estimates for the two SUBCLINICAL animals fall within or close to the range seen for CLINICAL suggests consistency in early phase titer kinetics, regardless of the initial disease severity, and supports the assumption that our observations captured the end of the initial stage of infection for these SUBCLINICAL animals. Furthermore, our findings are qualitatively and quantitatively consistent with a pattern of initial rapid antibody decay followed by a slower decay, as seen in other systems where long-term antibody titer kinetics were tracked within individuals [29, 30].

**Table 2.** Antibody titer decline rates and half-life values in days with their corresponding 95% confidence intervals [95% CI]. Data are reported for each individual in the CLINICAL and SUB datasets as well as for the first and second phase of titer decline observed for the SUB animals. Rates for the CLINICAL animals are ordered from high to low with the median decline and half-life values in ***bold italics***. The titer decline rate marked with an asterisk (*) was not significantly different from zero.

| Antibody Titer |  |  |  |
| --- | --- | --- | --- |
|  | Animal ID | Decline Rate | Half-life [95% CI] |
| CLINICAL | 1 | -0.156 | 6.4 [6, 6.9] |
|  | 2 | -0.149 | 6.7 [4.4, 14] |
|  | 3 | -0.099 | 10.1 [8.7, 12] |
|  | 4 | -0.075 | 13.4 [9.8, 21.4] |
|  | 5 | -0.065 | 15.5 [11.1, 25.9] |
|  | 6 | -0.061 | 16.3 [12, 25.3] |
|  | 7 | ***-0.058*** | ***17.1 [11.5, 33.3]*** |
|  | 8 | -0.058 | 17.2 [15, 20.2] |
|  | 9 | -0.058* | 17.3 [7.3, infinity] |
|  | 10 | -0.049 | 20.5 [13.6, 41.7] |
|  | 11 | -0.046 | 21.7 [13.1, 64.2] |
|  | 12 | -0.043 | 23.2 [19.2, 29.4] |
|  | 13 | -0.034 | 29.4 [16.1, 168.4] |
| SUB | 1 - 1st Phase | -0.037 | 26.8 [21.2, 36.6] |
|  | 1 - 2nd Phase | -0.001 | 975.9 [546.1, 4584.1] |
|  | 2 - 1st Phase | -0.164 | 6.1 [4.1, 12] |
|  | 2 - 2nd Phase | -0.002 | 433.4 [327.2, 641.4] |

To better understand the relationship between antibody titer and renal index, and to visualize how these biomarkers change relative to each other through time, we plotted the measures against each other to create a map of the host-pathogen space (Fig 3). With increasing time since infection, the CLINICAL animals followed a clear temporal trajectory, tracing a curved path starting in the high renal index and high titer space, dropping rapidly into the low renal index space with clinical recovery, and staying within the healthy range as antibody titers continued to drop (Fig 3A). In these CLINICAL animals, initial renal index values declined rapidly relative to antibody titers, so that only the earliest data points (<14 days since admission to rehabilitation) were found in the high titer, high renal index space. After 28 days, renal index scores leveled off within the healthy range and the temporal signal was dominated by antibody titer decline. By contrast, the SUBCLINICAL animals followed a straight path, always within the healthy range, as their antibody titers declined systematically throughout the 3 years that they were monitored (Fig 3A). All initial titers were high (log_2_ titer range CLINICAL=10-13, SUBCLINICAL=7-10) with variation among individuals observed. CLINICAL animals provided detailed information on initial changes in disease biomarkers, yet were released back into the wild within 6 – 12 weeks, providing no long-term data. In addition, as these animals stranded some unknown number of days after initial infection, the ‘upswing’ of antibody titers and clinical disease were not captured. Conversely, the SUBCLINICAL animals were followed for 3 years, providing important long-term biomarker data, but little on their initial dynamics (Table 1). Ultimately the CLINICAL and SUBCLINICAL paths overlapped, demonstrating convergence of the two trajectories and, potentially, similar long-term dynamics.

**Fig 3.**
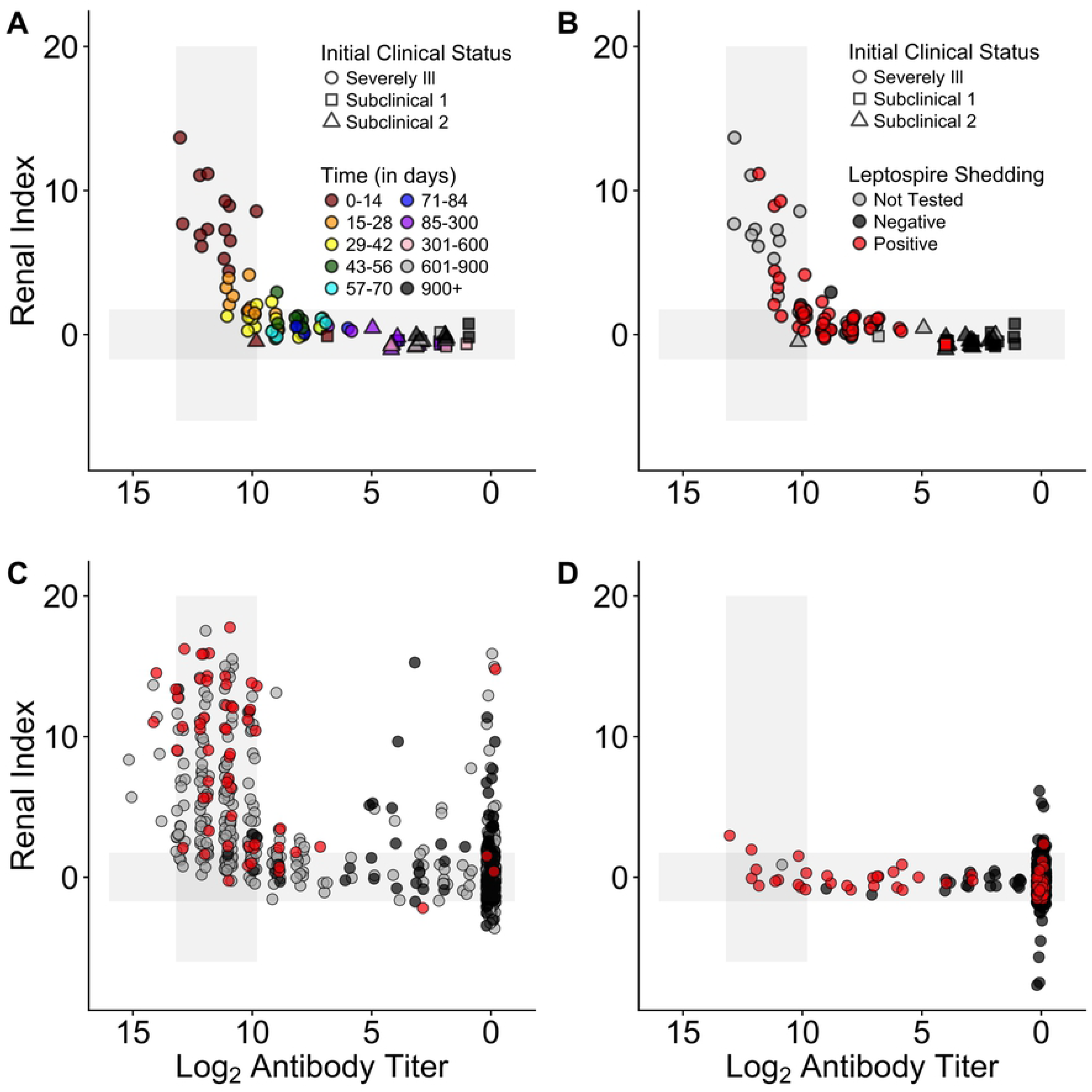
Map of host-pathogen space. Maps of the host-pathogen space created by plotting jittered log_2_ anti-*Leptospira* antibody titers (x-axis) against renal index values (y-axis). Plots created using data from the longitudinally followed animals (CLINICAL and SUBCLINICAL), color-coded by time since admission to a rehabilitation center (A) and by urinary leptospire shedding status (B). Plots created using cross-sectional data from stranded animals (STRAND; C) and wild-caught, free-ranging animals (WILD; D) color-coded by urinary leptospire shedding status. In all panels, horizontal grey bands are equivalent to those in Fig1 C&D, and the vertical grey bands are equivalent to the horizontal bands described in Fig1 A&B.

We used PCR to detect *Leptospira* DNA shed in the urine – a measure of current infection and potential transmission risk to others – and added pathogen shedding data to the map of the host-pathogen space. Addition of this third disease biomarker revealed that many animals continued to shed despite a rapid return to healthy renal function and systematic antibody titer decline (Fig 3B). All CLINICAL sea lions tested positive at least once in the first 38 days, most (11/13) continued shedding despite concurrent antibody titer decline and clinical recovery, and most (10/13) were still shedding at the last sampling point 4 – 12 weeks after initial admission (Fig 3B; also see [28]). Subclinical shedding of at least 8 weeks was detected in SUB1 [27], indicating that initial severe clinical disease is not a necessary condition for shedding of this duration. Shedding was never detected in SUB2, but the first urine testing date was 38 weeks after first detection of serum antibodies.

Altogether, our findings suggest that antibody titers act as a rough clock indicating time since exposure to the pathogen, with data on disease severity and pathogen shedding improving the temporal resolution of the host-pathogen trajectory.

### Using the Host-Pathogen Trajectory to Interpret Cross-sectional Data

Having established a temporal host-pathogen trajectory, we used our mapping approach to maximize the information gained from individuals observed only once. These cross-sectional data were from stranded (STRAND) and wild-caught, free-ranging (WILD) California sea lions (Table 1). When mapped, STRAND data fell along the trajectory mapped by the longitudinal data, but with greater variation, i.e., they cut a broader path through the host-pathogen space, and their map contained some outliers (Fig 3C). The STRAND data contained a wider range of renal index scores (−3.6 – 17.8) and a higher maximum antibody titer (log_2_ titer = 15) than did the longitudinal data (renal index = −1.0 – 13.7; maximum log_2_ antibody titer = 13; Fig 3A-C), suggesting that STRAND data captured a greater overall range of sea lion-*Leptospira* host-pathogen dynamics than the smaller dataset of longitudinally followed animals. The larger size (50-fold larger than CLINICAL) and broader selection conditions (i.e. including animals so ill from leptospirosis they died quickly, as well as those compromised for other reasons) of the STRAND dataset could explain this difference. By contrast, and in keeping with our assessment of apparent health at capture, the WILD animals chiefly occupied the space defined by the SUBCLINICAL and the recovered CLINICAL animals (Fig 3D). Notably, the WILD and STRAND datasets both differed from the SUBCLINICAL and CLINICAL in that they contained substantial numbers of seronegative animals.

We analyzed the distribution of renal index scores in each group, using antibody titer levels to standardize for time since infection, in order to assess (1) whether each group had a unique renal index profile or whether the SUBCLINICAL, WILD and CLINICAL groups were merely opposite extremes within the range seen in the STRAND animals, with WILD and SUBCLINICAL at one extreme and CLINICAL at the other, and (2) whether all groups converged to the same point with time since infection (Fig 4; Table 3). Renal index distributions of WILD and SUBCLINICAL animals never differed significantly. Those of the WILD and CLINICAL animals differed significantly at each antibody titer level assessed, yet the difference between their mean renal index values decreased as titers decreased, i.e. they were converging with time since infection. While some STRAND animals exhibited markedly greater renal disease than WILD animals, at all titer levels, there is also substantial overlap between these groups, suggesting that the WILD animals are similar to the majority of STRAND animals not suffering from clinical leptospirosis. Altogether, for all datasets and regardless of the starting point in the trajectory (i.e. the renal index value at the highest titers), as antibody titers declined, so did mean renal index scores and the trajectories of each of the different datasets converged towards the healthy range.

**Fig 4.**
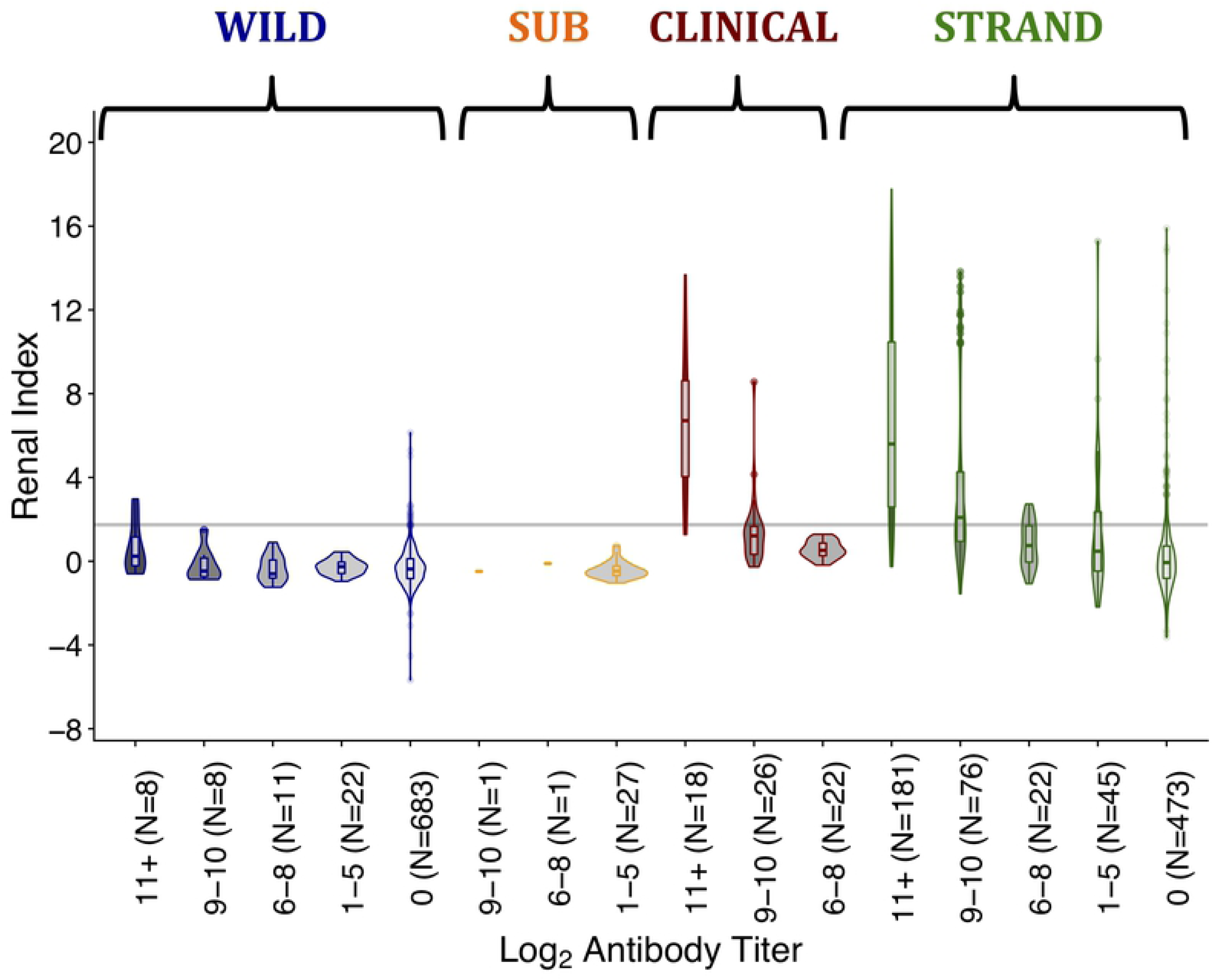
Renal index score distributions by log_2_ antibody titer level, for each sample group. Sample groups are as described in the methods and are wild-caught (WILD), subclinical (SUB), clinical (CLINICAL), stranded (STRAND) sea lions. Titer groups were chosen based on the titer dynamics observed in the longitudinally followed animals through time. Groups roughly match the different phases of the host-pathogen relationship ranging from initial infection (11+), clinical recovery (9-10) and two stages of historic infection (1-5 and 6-8). Group 0 contains seronegative animals. The grey line denotes the upper boundary of the healthy range for renal index values. The upper whisker extends from the hinge to the highest value that is within 1.5 * IQR of the hinge, where IQR is the interquartile range. The lower whisker extends from the hinge to the lowest value within 1.5 * IQR of the hinge.

**Table 3.**
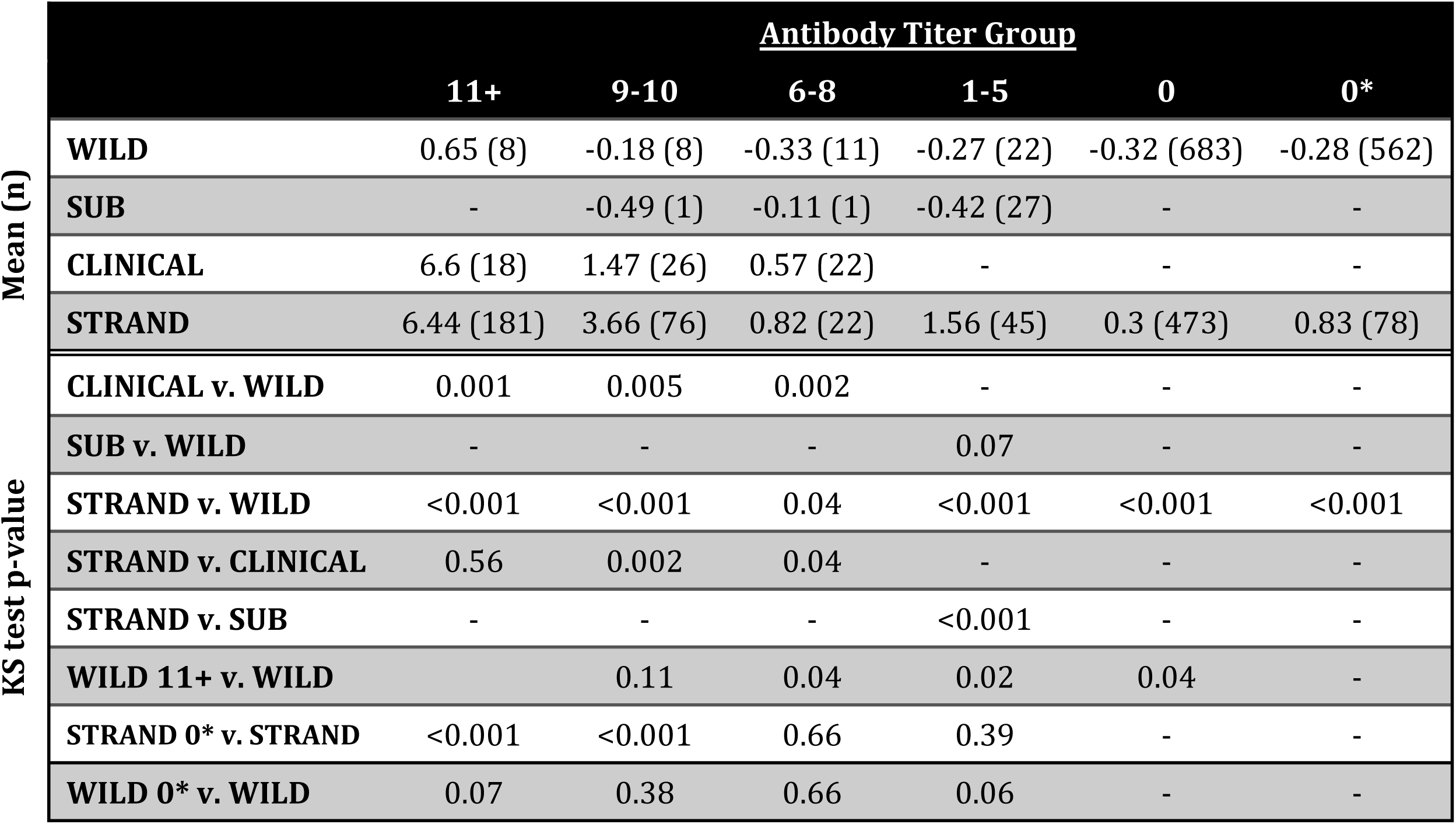
Mean renal index scores and sample sizes (n) by antibody titer group for initially clinical (CLINICAL), initially subclinical (SUB), stranded (STRAND) and wild-caught (WILD) animals. Titer group “0” contains all seronegative animals, titer group 0* contains only non-shedding (i.e. urine PCR negative) seronegative animals. P-values are for two-sided bootstrap Kolmogorov-Smirnov (KS) test comparisons between animal groups of renal index distributions for a given titer. P-values for WILD 11+ v. WILD are for one-sided bootstrap KS test comparisons within the WILD dataset, with the null hypothesis that the renal index distribution for the 11+ titer group will be greater. P-values for STRAND 0* v. STRAND are for two-sided bootstrap KS test comparisons within the STRAND dataset comparing renal index distributions of seronegative, non-shedding animals with the other titer groups.

As with the longitudinally sampled animals, leptospiral DNA was detected in both STRAND and WILD animals for a wide range of antibody titer and renal index values (Fig 3C&D). However, unlike the longitudinal groups, the cross-sectional data also included seronegative animals (i.e. no detectable anti-*Leptospira* antibodies; Fig 3C&D; plotted above log_2_ titer of 0). These animals presented with a range of renal index scores and, intriguingly, included animals shedding leptospiral DNA (Fig 3C and 3D; see section ‘Antibody Titer Kinetics and Shedding Duration’ for further discussion of these animals).

The broad congruence of the cross-sectional datasets with the longitudinally collected data corroborates the assumption that the CLINICAL and SUBCLINICAL animals jointly define the course of infection in this space and establishes that cross-sectional data can be interpreted within this temporal framework.

#### Tracking the Distribution of Disease Severity

Defining the mean and range of pathogen-induced disease severity at different times since infection enhances our ability to interpret confusing host presentations (Fig 2) and hence understand disease dynamics in a system. However, biases in data sources must be considered when interpreting these data. In our study, when all data are combined, we see that initial disease severity (i.e. renal indices when animals have high antibody titers) ranges from healthy to severely ill (Fig 3). However, by design, initial renal index values in the CLINICAL animals captured only the upper range of disease severity, while the SUBCLINICAL animals occupied only the lower healthy range. WILD animals were sampled only if apparently healthy, and their renal index scores reflected this initial assessment, mostly occupying only the healthy range even during the presumed initial stage of infection (Fig 3 and Fig 4 titer level 11+). By contrast, STRAND data, which were collected without applying selection criteria to candidate animals, showed a wide range of initial disease severity and appear to knit together the various subset datasets to which specific selection criteria were applied (e.g. WILD, CLINICAL, SUBCLINICAL; Fig 3). Of note, although most of the seropositive WILD animals fall within the healthy range of renal index values, at the highest titer values a few exceed the healthy range (Fig 3D), and the mean renal index score of those individuals at this highest titer level is greater than those of the other levels (Fig 4; Table 3), suggesting that these animals can experience some degree of initial renal compromise from which they recover.

In many systems, disentangling disease caused by the pathogen of interest versus disease from another etiology can be difficult. In our study, while STRAND data capture the full spectrum of disease and follow the trajectory defined by the longitudinal data, this trajectory is shifted up the renal index axis and there are some obvious outliers (e.g. mid-low antibody titer, high renal index individuals; Fig 3C). STRAND renal index score distributions were significantly higher than those of almost all other datasets (i.e. CLINICAL, SUBCLINICAL, and WILD) at all antibody titer levels (with the single exception that renal index distributions for the highest-titer groups of STRAND and CLINICAL were indistinguishable (Fig 4; Table 3)). We hypothesize that this upward shift in STRAND renal index score is due to individuals experiencing renal compromise from causes other than leptospirosis and that overall STRAND host-pathogen dynamics are consistent with those described by the longitudinal data, i.e., animals are recovered clinically from leptospirosis by the time their log_2_ antibody titers have declined below 9 (Fig 3A-C & Fig 4).

To test this idea, we analyzed the group of seronegative, non-shedding STRAND animals that were presumably never infected and never exposed. Any renal compromise observed in this group would be from a cause other than leptospirosis and the range of their renal index values provides a reference against which to compare currently or previously infected sea lions. We found that the renal index distribution of these seronegative, non-shedders in the STRAND dataset (denoted 0* in Table 3) was not significantly different from those of the mid and lower antibody titer STRAND groups (1-5, 6-8; (Fig 4; Table 3). This suggests that outliers found on the map – mid-low antibody titer with high renal index (Fig 3C) – which, according to the host-pathogen trajectory described in Fig 3A-B, should have fully recovered from *Leptospira*-induced renal compromise, are equivalent to the seronegative non-shedding STRAND animals experiencing disease from another etiology. Similar analyses of the WILD animals showed no significant difference between seronegative, non-shedding animals and animals with titers, further supporting our assumption of apparent health of this group.

#### Predicting Survival

In hospital and rehabilitation settings, determining probability of survival can be vital for patient triage and efficient allocation of resources. Using data on renal index scores that are readily available at admission, we found a significant negative relationship between these scores and survival of animals suspected of having leptospirosis (Fig 5; OR = 0.64, 95% CI = 0.53 – 0.78, p<0.001). This relationship is not only informative for guiding management in a clinical setting, but the absence of high renal index values in the WILD animals suggests that *Leptospira*-associated mortality in the apparently healthy animals selected for sampling is likely low.

**Fig 5.**
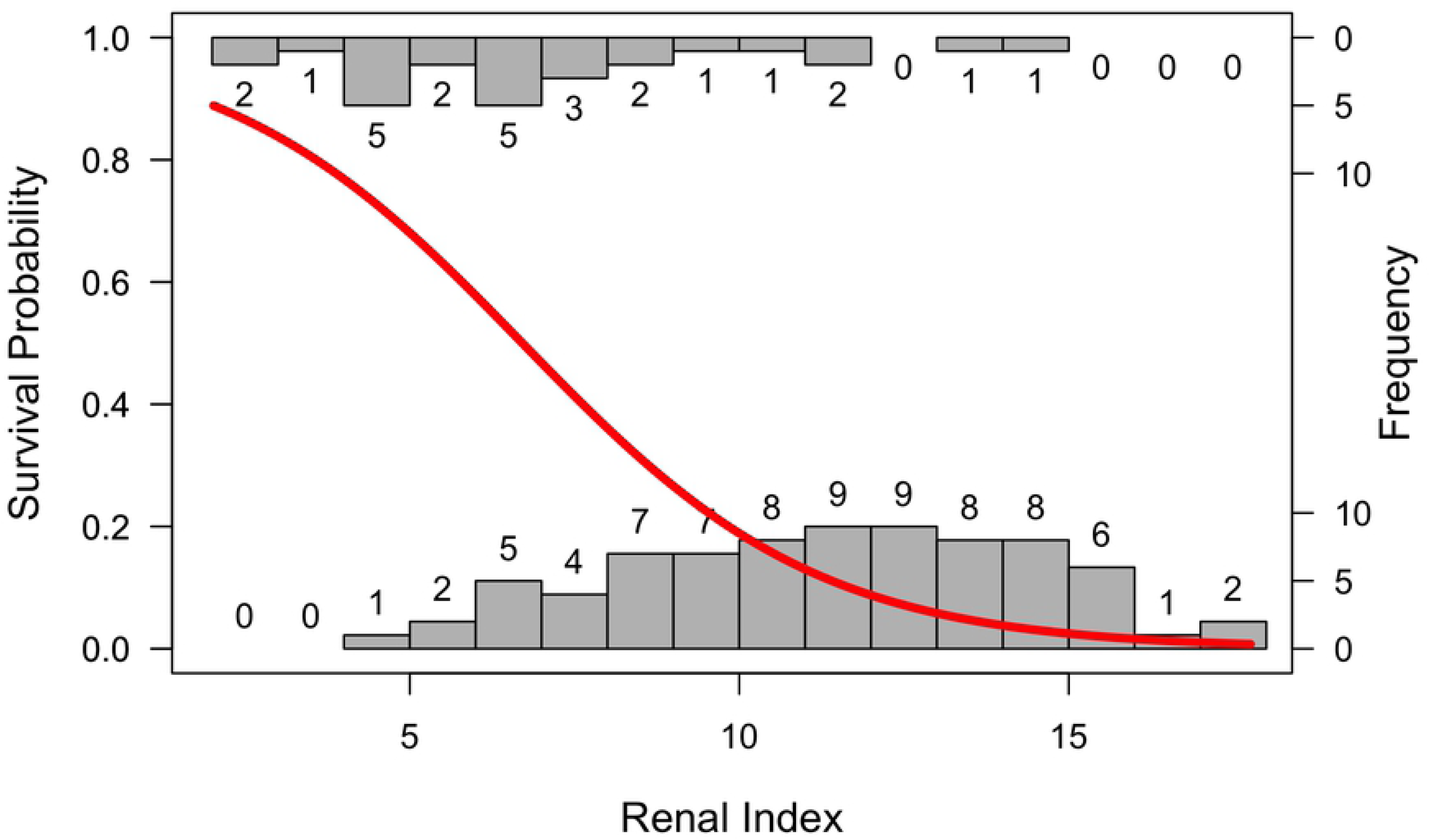
Survival probability predicted from renal index score. The probability of survival as a function of renal index score plotted over histograms of the number of animals surviving (top histogram) or not surviving (bottom histogram) by renal index value. Analyses were run using data from samples collected within 72 hours of admission to a rehabilitation center from animals stranding and diagnosed with leptospirosis.

#### Antibody Titer Kinetics and Shedding Duration

Estimating an individual’s time since infection aids in assessing infection incidence [6, 16] [4] and, in combination with data on shedding status, can enable estimation of the duration of infectivity – a value which is notoriously difficult to determine in wildlife where repeated sampling of individuals is rare. To approach this problem, we explore the hypothesis that for our system there is a single dominant pattern of antibody titer decline, regardless of initial disease severity, such that titer acts as a rough measure of time since infection. We begin by noting that animals shedding leptospiral DNA exhibited antibody titers ranging from very high to seronegative (Fig 3B-D; S2 Table). Under the working hypothesis that all animals experience similar antibody titer kinetics, low titer and seronegative shedders would be chronic shedders.

To test this idea we considered the epidemiological context of our data: during our study period, outbreaks occurred in 2008 and 2011 (Fig 6A). If antibody titer decline initially occurs rapidly as seen with the CLINICAL animals and then quite slowly as seen with the SUBCLINICAL animals (Fig 2C&D), we should see relatively few low titer animals during an outbreak year, and relatively more with each year until the next major outbreak. Similarly, observation of seronegative shedders should occur after outbreaks as titers dip below detection.

**Figure 6.**
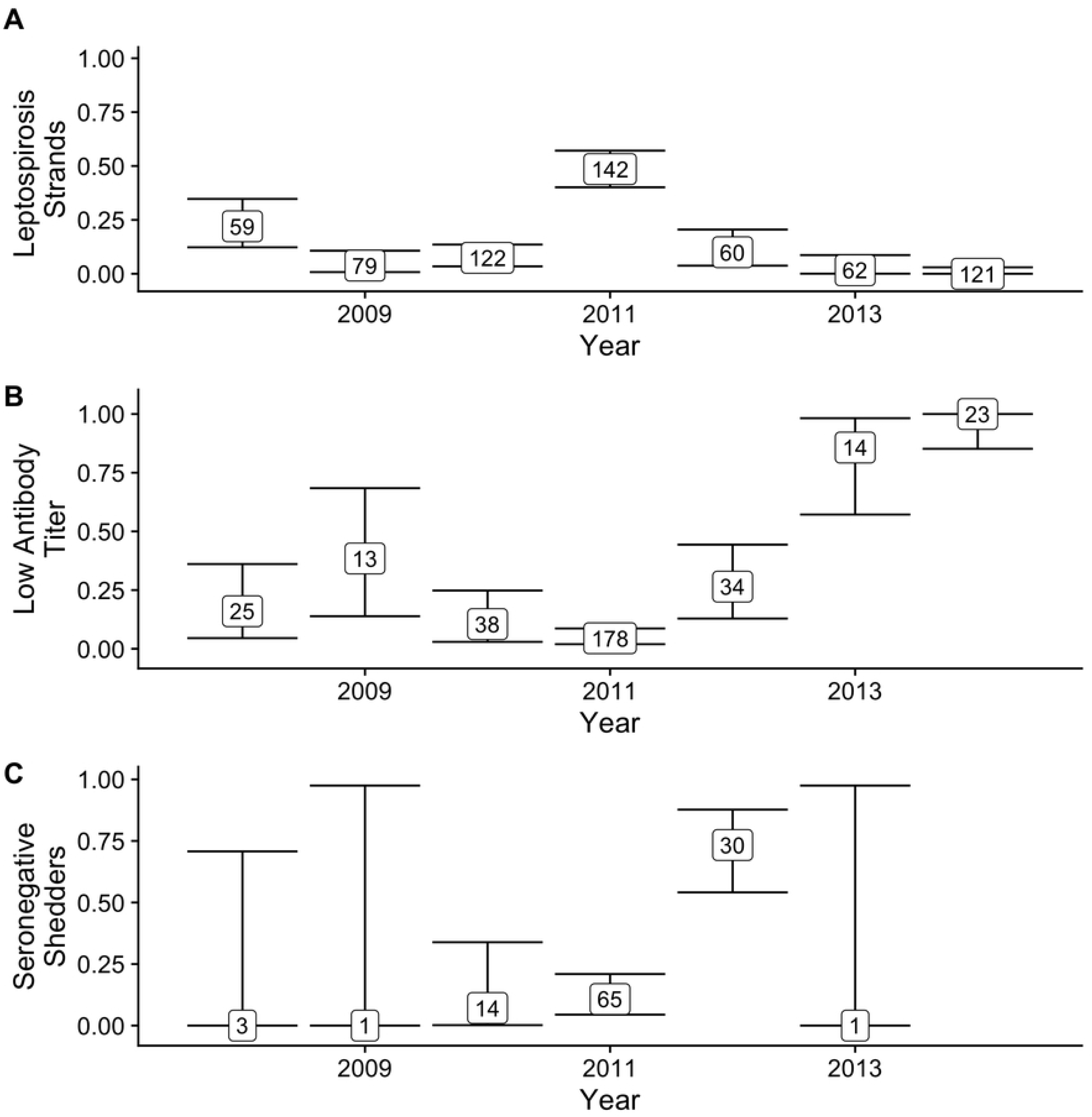
Proportion of stranded animals with leptospirosis (A), proportion of seropositive animals that have low titers (log_2_ titer 1-5) by year with 95% CI (B), and proportion of shedding animals that are seronegative by year with 95% CI (C). Total sample size for each proportion is indicated within the box. Only STRAND data included in (A) WILD and STRAND data for (B) and (C). The proportion of leptospirosis strands is highest in the two outbreak years – 2008 and 2011 – and the proportion of low titer animals increases with each year after the outbreaks. Similarly, the proportion of seronegative shedders increases after the major outbreak in 2011, but then declines to zero by 2013. The single shedder in 2013 had a log_2_ antibody titer of 3, no animals were shedding in 2014; therefore a proportion could not be calculated. Few shedders were detected in 2008 and 2009 due to small sample sizes of animals PCR tested for shedding (2008 N=6, 2009 N=3, 2010 N=73, 2011 N=158, 2012 N=119, 2013 N=162, 2014 N=291).

We found support for our hypothesis when we plotted data from the combined WILD and STRAND datasets. The proportion of low titer (log_2_ titer 1-5) animals increased following major outbreaks, particularly after 2011 when incidence remained very low (Fig 6A&B). Regarding seronegative shedding, as predicted the proportion of shedding animals that were seronegative was high in 2012, immediately following the 2011 outbreak; however, intriguingly, no seronegative shedders were detected in 2013 or 2014 (Fig 6C). No seronegative shedding was detected in 2008 and 2009 either, but this is likely due to very small sample sizes of animals tested in these years (N=3-6) compared with 2010-2014 (N=73-291; S2 Table). The fact that seronegative shedding occurred the year immediately after a major outbreak, but not in the two years that followed, suggests that titer decline may occur more rapidly in some individuals than predicted by SUBCLINICAL animal titer kinetics and that shedding duration may be less than 2 years. In addition, the prevalence of longer-term chronic shedders might be quite low, resulting in low power to detect.

Although noisy, these field data align with our argument that animals that are shedding while seronegative (or low-titer seropositive) may be chronic shedders. Combined with our earlier result that antibody titers act as a rough measure of time since infection, this provides an opportunity to learn more about shedding duration. Precise quantitative estimates are impossible, particularly due to wide uncertainty on the slow decay rate of low titers, but a lower bound on shedding duration can be computed using the initial rapid decay rate. Assuming a constant titer decay rate of 0.058 log_2_ antibody titer units/day (the median decay rate of the CLINICAL animals) and an initial titer of 11 (the median initial titer of the CLINICAL animals), we conservatively estimate the approximate time taken to reach a given titer level (S3 Table), e.g. we calculate that it takes at least 189 days to reach seronegative status. From this, we can estimate the approximate duration of shedding for a PCR-positive individual with that antibody titer. However, applying similar logic to the decay curves suggested by the SUBCLINICAL animals, and still assuming an initial titer of 11, we obtain estimates of shedding duration for seronegative shedders that are much longer, e.g. ∼ 6 years. Given the important caveats stated above and the low SUBCLINICAL sample size (N=2), as well as further biological caveats discussed below, the true shedding duration of seronegative shedders likely falls somewhere between these two estimates.

#### Predicting Shedding

Data on antibody titers (and in our case renal index, also derived from serum samples) are often more readily available than those on active pathogen shedding. If a clear relationship between one or both of these biomarkers and shedding can be established, then shedding may be predicted when shedding data are otherwise unavailable. Using a dataset including all animals (CLINICAL, SUBCLINICAL, WILD and STRAND) for which shedding data were available, we performed logistic regression to investigate how antibody titer and renal index were related to pathogen shedding (using only the first sample date per individual). We found that antibody titer contributed significantly to the final model (p<0.001), but that renal index (p=0.96) and the interaction between renal index and antibody titer (p=0.85) did not. The probability of shedding increased with increased antibody titer (Fig 7A; OR = 1.67, CI = 1.55 – 1.80). Using this relationship, we predicted shedding status in STRAND animals for which shedding data were missing, using only the first sample date per individual, and were able to produce a more complete map of the host-pathogen space (Fig 7B&C). Using all datasets, but including only the first sample collected for an animal, we estimated an overall shedding prevalence of 0.22 (PCR tested and untested, n=1510) which is substantially higher than the prevalence of 0.15 calculated using the raw data (PCR tested only, n=811), showing we may be greatly underestimating shedding prevalence when shedding data are rare relative to antibody titer data.

**Fig 7.**
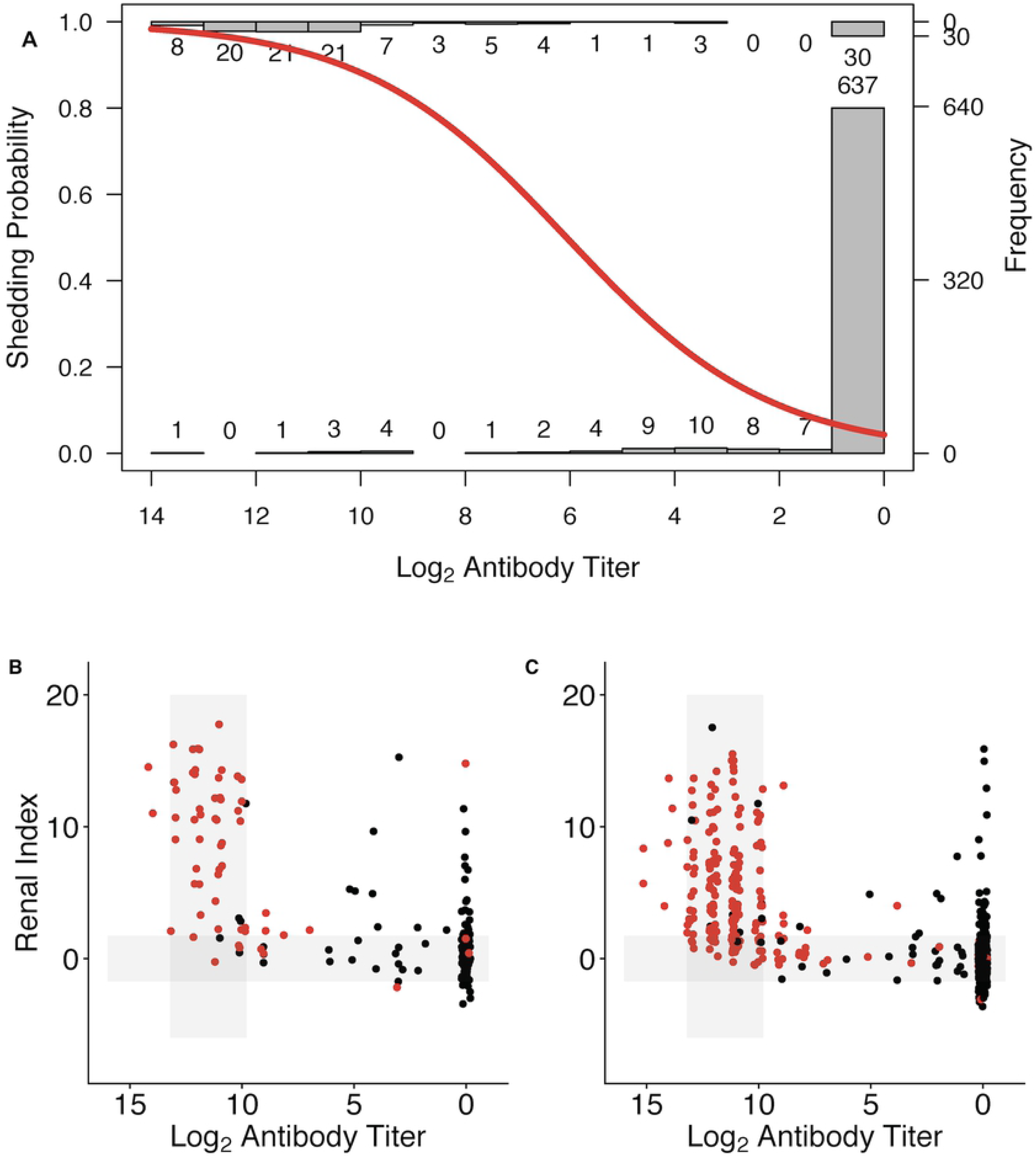
Shedding probability predicted from log_2_ anti-*Leptospira* antibody titer. (A) The probability of shedding as a function of log_2_ anti-*Leptospira* antibody titer plotted over histograms of the number of animals shedding (top histogram) and not shedding (bottom histogram) by antibody titer (A). STRAND data plotted using the ‘host-pathogen map’ framework as in Fig 3C. Data divided into those individuals that were PCR tested and for which shedding status was known (B), and those that were not PCR tested and for which shedding status was predicted (C; positive = red, negative = black).

## Discussion

We have introduced a host-pathogen mapping framework that characterizes the progression of *L. interrogans* infections and clinical responses in California sea lions, drawing on longitudinal data from individual animals. The usefulness of our host-pathogen map for interpreting cross-sectional data arises from the overall consistency in biomarker dynamics across individuals, and particularly within similar groups of individuals. In longitudinally sampled animals, we found antibody titer acted as a rough clock marking time since infection. Although there was heterogeneity in initial antibody titers and decline rates, animals followed the same broad initial pattern of titer decline regardless of pathogen shedding status and initial disease severity. Our longitudinal data were censored – we lacked data from the earliest stages of infection in these stranded animals, especially for the SUBCLINICAL animals, and the CLINICAL animals were followed for at most 90 days – yet trajectories of antibody titer decline for both groups overlapped and converged, suggesting that ultimately they follow the same long-term dynamics. Analysis of cross-sectional data corroborated this finding, as they traced the same broad trajectory as the overlapping SUBCLINICAL and CLINICAL groups, knitting them together. In addition, consistent with our observation of an initial rapid antibody decline followed by a slow second phase of decline, cross-sectional data revealed that in the years following a major outbreak and before another one occurred, the relative proportion of low titer animals increased with time as initially high titers followed biphasic decline kinetics.

Our host-pathogen map, which shows how severely ill and subclinical infections are linked, enables us to map the complexity of the host-pathogen relationship, resolve questions about apparently anomalous presentations, and is useful in addressing a particular challenge with respect to the traditional dichotomous view of *Leptospira*-host relationships. This view describes host species as either reservoir hosts that experience little disease but can shed chronically for months to years, or accidental hosts that can become severely ill, and possibly die, but do not become chronically infected [1, 25]. Thus, sea lions present an interesting challenge to this view as they show characteristics of both presentations. For example, sea lions showed a range of clinical disease in the initial stage of infection (i.e. at high titers), and although shedding was detected in the earliest phase regardless of clinical disease, there was substantial variation in shedding duration as determined by detection of both shedding and non-shedding sea lions at each antibody titer magnitude (including seronegative). Our mapping approach resolves this tension by showing that, in the case of *Leptospira* infection in sea lions, the accidental and reservoir characteristics seen in individual sea lion presentations are extremes of a unifying trajectory of the host-pathogen interaction, and our map shows how these classical manifestations are linked both within an individual’s infection as well as on a population level. This approach is of particular value for wildlife disease ecology, since many host-pathogen relationships are poorly characterized, and an in-depth study in a controlled experimental setting is generally not possible.

Characterizing the temporal trajectory of the infection and recovery process and establishing consistency in antibody kinetics is especially important for accurately interpreting the relationship between antibody titer and leptospiral DNA shedding. This enables rough estimation of shedding duration and potentially identification of chronic shedders – data that are key to accurate model parameterization, but notoriously difficult to collect for wildlife systems. Infectious disease theory predicts that shedding duration will influence transmission dynamics and modeling efforts have shown that chronically shedding individuals play a critical role in population-level pathogen persistence both in general [31] and specifically in California sea lions [19]. Using our map of titer decay and clinical recovery, we identify low titer and seronegative shedders as likely chronic shedders, and using titer kinetics of longitudinally followed animals we obtain estimates of shedding duration from animals sampled only once.

Quantitative analysis of biomarker data can also define relationships between biomarkers and specific disease outcomes. Results from these analyses can then be applied to fill gaps in incomplete datasets. This is particularly important as some desired data types are more difficult to obtain, or are unavailable at particular time points, e.g., survival data are only available when no longer clinically useful, and urine can be more difficult to collect than serum. We predicted urinary shedding of *Leptospira* from antibody titer magnitude, and by applying this relationship across our full dataset we estimated a higher shedding prevalence than that in the smaller dataset for which shedding data were available. However, the opposite could have been true had the group composition been different in the smaller ‘training’ dataset. Therefore, in all cases of establishing these relationships between biomarkers, the impact of group composition and the epidemiological context of the data must be carefully considered. For example, in our case, seronegative shedding was more common the year following a major outbreak, had only these data been used when establishing the relationship between titer and shedding, shedding prevalence estimates would have been much higher. In future work, increasing the amount of data included – biomarker, demographic, environmental – in models defining these relationships, as in Borremans et al. [6], may further improve estimates by accounting for differences among individuals, and time periods.

By analyzing quantitative antibody titers jointly with other biomarker data, such as clinical or infection status, individuals that are indistinguishable by one measure but in fact are biologically distinct, may be better characterized and identified. For example, subclinical infection was seen across all titer levels, but severe disease was seen almost exclusively at high titers – as titers declined, only the rare outlier showed evidence of clinical disease. Using a binary approach to interpreting antibody titers, the outlier individuals with mid-low titer and severe renal disease may be categorized as ‘seropositive’ and miscategorized as acute *Leptospira* infections. However, through comparisons with a group of individuals thought never to have been infected, we show that these outliers were infected in the past (giving rise to the detected antibodies) but likely exhibited renal disease from another more recent etiology. This argument is supported by the lack of leptospiral DNA shedding in any of these outlier individuals. This general principle is relevant to the broader field of public health, as a pervasive health issue in developing countries is the similarity in disease presentations among a diverse group of infectious and non-infectious etiologies (e.g. acute febrile illnesses; pneumonia; diarrheal disease [32, 33]) leading to possible misdiagnosis. Quantitative analysis of multiple biomarkers, as exemplified by our host-pathogen map, facilitates identification of outliers and thus cryptic causes of disease. This information can aid appropriate treatment choice by clinicians, management recommendations by epidemiologists, and accurate estimation of health and economic burden of a particular pathogen by public health agencies [34, 35].

Using our map and analyses of multiple biomarkers we are able to make predictions regarding survival, shedding duration, and etiology of clinical disease. We propose that our approach, or a similar one, may be applied to other host-pathogen systems, but system-specific modifications may be required. In some systems, antibody titer dynamics may contain more heterogeneity than seen in the *Leptospira*-sea lion system [6, 15, 16, 20, 21], necessitating adjustments in the construction or interpretation of the host-pathogen map such as using different biomarkers, or including more of them [3, 6]. For example, working in an experimental system, Torres et al. [3] used blood pathogen concentration, instead of antibody titer, and multiple measures of host health to build a map of ‘disease space’ within which individuals that became severely ill and died, and those that survived, traced different pathways as they moved from time since infection. They hypothesized that using this map they could plot cross-sectional data and infer an individual’s infection time-line and predict their prognosis, but did not test this idea.

Heterogeneity in the maximum antibody titer, degree of clinical disease experienced, the shape of the titer decline curve, and the duration of detectable antibodies has been noted in a number of host-pathogen systems [8, 10, 13, 15, 20, 21], and in some cases, such heterogeneity is associated with specific characteristics of an individual’s infection. Subclinical infections with other pathogens have been associated with lower maximum antibody titers [15, 21], and with shorter antibody titer persistence [15]. Therefore, it was not obvious *a priori* whether antibody kinetics of subclinical and clinical infections would be the same in California sea lions. However, our findings, and those of others, indicate that despite some heterogeneity in antibody titer magnitude, titer kinetics for clinical and subclinical infections were roughly the same for *Leptospira* in sea lions and Q fever in humans [22].

Similarly other studies have examined whether chronic infections might be associated with different titer dynamics, e.g. higher maximum titers and longer persistence have been seen for chronic cases of Q fever [13, 20]. However, if anything, our data show the opposite trend. Instead of higher titers and greater antibody persistence, some of our hypothesized chronic carriers had no detectable antibodies. While our two subclinical animals – neither of which shed beyond the first several months of infection – had detectable antibody titers for years. Long-term persistence of antibodies in the SUBCLINICAL animals may be due to their captive status and its impact on overall condition and immune function. Alternatively, the long duration of detectable antibodies in the SUBCLINICAL animals and the lack of detectable titers in some of our shedders may reflect the full range of expected variation in titer decay rates and hence titer persistence.

The antibody titer decline that we detected in our sea lions, despite continued infection in some, may be due to pathogen-specific differences in the underlying host-pathogen interaction. For example, with some infections, including Q fever, the pathogen continues to circulate in the blood in chronically infected individuals, [36], stimulating the immune system to continue to produce antibodies and resulting in persistently high titers. Conversely, in chronic *Leptospira* infections, once leptospires have colonized the kidneys they appear to evade the host immune system [37, 38], which would explain the observed antibody titer decline in our system, despite chronic renal infection and shedding.

We believe our host-pathogen mapping approach yields many benefits, however the following caveats – some specific to our system, some more generalizable – must be considered. Individuals in any study population will experience differences in environmental exposures and conditions and we know that this, and age specific differences, can lead to variation in biomarker data. For example, adaptive immunity can be influenced by many factors including age, nutritional status and pathogen exposure history [39–41]. In addition, several idiosyncrasies in our study may have affected our findings. Estimates from the two SUBCLINICAL animals may not precisely reflect population level trends given the small sample size of the SUBCLINICAL group. Our observations were censored, as data from the CLINICAL animals were limited to the early phase of disease and recovery, there were only a few data points from this early phase of infection in the SUBCLINICAL animals, and the very earliest phase from infection to initial illness was unobserved for all animals. The initial infecting dose of *Leptospira* is unknown in all cases and may have varied substantially, potentially impacting immune response and disease severity. Both SUBCLINICAL and CLINICAL animals experienced potentially immunomodulatory conditions, specific to their time in captivity, that animals in the wild would not have. For example, the SUBCLINICAL animals remained in captivity where they were neutered and maintained in excellent body condition in a controlled, predator-free environment. Such conditions may have increased their reserves and their capacity to invest in a costly immune response [42], resulting in differences in long-term antibody titer kinetics relative to free-ranging animals, i.e. a slower decline and more persistent antibody titers. Similarly, CLINICAL animals received medical treatment and supportive care which may have affected survival, increased their reserves and boosted their immune potential. Together, these factors may help explain why estimates of shedding duration for seronegative shedders, based on SUBCLINICAL antibody titer kinetics (i.e. roughly 6 years to become seronegative), differed from patterns of seronegative shedding detected in the wild after a major outbreak (i.e. decline to seronegative within 2 years). However, despite the unique circumstances of our SUBCLINICAL and CLINICAL animals, the combined longitudinal datasets describe a multiphase titer decline consistent with that found in other studies [29, 43], and overall patterns seen in the longitudinal data were consistent with those in the cross-sectional data and likely reflective of the entire sea lion population.

Approaches that integrate biomarker kinetics to interpret cross-sectional data can be useful to clinicians and ecologists alike, and bridge perspectives from these often separate worlds. Clinicians tend to focus on individual health, while ecologists focus more on quantifying the natural process and understanding disease dynamics at the population scale. Using this integrated approach, clinicians can improve individual patient survival through more accurate patient triage and efficient allocation of resources, and can reduce transmission risk to others. Treatments that are expensive, of limited availability, or time intensive (e.g. dialysis) may be reserved for those individuals with the most severe disease and the lowest probability of survival in the absence of such therapy. Conversely, in a wildlife rehabilitation setting, limited resources might be directed towards those with the highest probability of survival. Stricter, but possibly more expensive, measures to prevent transmission can be efficiently directed at those individuals with the highest probability of shedding. Ecologists can better conceptualize model structure using estimates of shedding prevalence and duration and can better describe population level transmission dynamics. For example, Buhnerkempe et al. [19] found that the addition of a chronic shedder compartment to the traditional SIR (susceptible, infected, recovered/resistant) model was necessary to accurately describe California sea lion-*Leptospira* dynamics and to capture long-term patterns of pathogen persistence. Once model structure has been determined, survival probabilities based on quantitative data (e.g. health, antibody titers) will influence the duration spent in various model compartments and thus how they contribute to onward transmission or herd immunity.

Historically, many of the principles of disease ecology were developed with childhood diseases such as measles in mind, and these acute infections have much crisper life histories for which the relatively simple SIR models can be used to capture the dynamics [44] (See Fig 1 and S1 Box). As the field addresses more complex host-pathogen relationships, these old assumptions break down and other models and approaches are needed. Models need to include greater complexity such as longer and variable infectious periods [19, 31], quantitative data and antibody titer kinetics [6, 15, 18], and multiple biomarkers of disease [6] [4]. Models based on quantitative data and that integrate antibody titer kinetics have been found to result in better estimates of model parameters (e.g. transmission rate, basic reproductive rate) and improved model performance and predictive capability (e.g. force of infection, incidence of infection), especially when only cross-sectional data were available [6, 10, 12–14, 16–18]. Similarly, models integrating quantitative serologic data may provide better estimates of incidence, especially relative to estimates based on reported rates of illness, as such reports miss subclinical infections [12] [6, 15–17]. We propose that our host-pathogen map provides a framework with which to visualize quantitative data from multiple biomarkers, determine the relationships between them, and identify the temporal trajectory of infection and recovery as reflected in changes in biomarker levels through time. This approach is especially useful for elucidating pathogen dynamics in wildlife systems, which typically rely on cross-sectional data. Ultimately this approach can clarify the biology of more complex host-pathogen systems, and enable the design of more appropriate dynamical models and statistical methods.

## Materials and Methods

### Study Animals

An overview can be found in Table 1.

#### Wild-caught California sea lions

Urine (n=637) and serum (n=732) samples were collected from anesthetized or physically restrained unique sea lions (n=730; 2 animals were recaptured and resampled) caught between September 2008 and November 2014 from three regions – southern California (San Miguel and San Nicolas Islands), central California (Año Nuevo Island, Monterey and San Francisco’s Pier 39) and northern Oregon (Astoria, OR). All urine collection occurred under anesthesia. To minimize anesthetic risk, only apparently healthy animals were captured and sampled. Estimated ages ranged from 1 to 5 years. These animals represent a cross-sectional sampling of the apparently healthy, wild, free-ranging population and we refer to them as “WILD”.

#### Stranded California sea lions

Urine (n=166) and serum (n=797) samples were collected from 724 unique California sea lions that stranded along the central and northern California coast and were admitted to a marine mammal rehabilitation center (The Marine Mammal Center, Sausalito, CA) between 2008 and 2014. Animals stranded due to illness or injury of all kinds, including, but not limited to leptospirosis, domoic acid toxicity, trauma, neoplasia, pneumonia and malnutrition. To match the age range for the wild-caught animals, only animals between the ages of 1 and 5 years were included in the study. These animals represent a cross-sectional sampling of the ill and injured sea lion population and we refer to them as “STRAND”. Only STRAND animals with both Microscopic Agglutination Testing (MAT) and chemistry results from within 14 days of each other were included in the study (>95% were from within 24-hours of each other). Urine PCR results were included only if these results were from urine collected within 24 hours of serum collection for chemistry analysis. This was to ensure that data on current infection (PCR) was from the same time point as data on clinical status (serum chemistry), which can change substantially quite rapidly. Serum antibody titers have a much slower rate of change, therefore we allowed serum MAT results to be within 14 days of PCR and serum chemistry sample collection dates.

#### CLINICAL

In 2010 and 2011, The Marine Mammal Center rescued and initiated treatment on 91 subadult, juvenile and yearling sea lions that stranded due to severe leptospirosis. Of these, 66 died, typically within days of stranding (median = 4 days, interquartile range =2-7). We tracked the progression of host response (serum chemistry values, anti-*Leptospira* antibody titers) and active infection (leptospiral DNA shedding in urine) in 13 sea lions that survived to be released, and we refer to them as “CLINICAL” animals. Animals were diagnosed with leptospirosis using a combination of clinical observation, serum chemistry data and necropsy data [45]. Animals were longitudinally sampled starting on Day 0 (their first day in rehabilitation; serum only) and then approximately every 14 days thereafter (serum and urine) until an individual’s release back into the wild 6-12 weeks later, as described in Prager et al. [28]. These animals were not included in analyses of the larger STRAND dataset except when specifically noted.

#### SUBCLINICAL

Two animals, which we will refer to individually as SUB1 and SUB2, collectively as SUBCLINICAL, never showed clinical signs of disease the entire period during which they were monitored as determined by clinical observation, complete physical examinations, complete blood counts and serum chemistry data. SUB1 stranded as a yearling male (i.e. between 1 and 2 years) in southern California in June 2010 (as described in Prager et al. [27]) with a flipper injury that precluded release back into the wild, and initially had no detectable anti-*Leptospira* antibodies. SUB1 seroconverted (i.e. acquired anti-*Leptospira* antibodies) in rehabilitation, with no observed clinical signs, at some unknown time-point within a 15-month period and was shedding leptospiral DNA 62 days after the first detected anti-*Leptospira* antibodies [27]. SUB1 was adopted by the U.S. Navy Marine Mammal Program (MMP) July 12, 2012, and samples were provided for monitoring every 1 to 9 months for a total of 44 months from the initial date that antibodies were detected. Because the date of infection was unknown, for our analyses we used the date of the first detected titer as Day 0. SUB2 stranded as a juvenile male (i.e. between 2-4 years), moderately underweight with a neck laceration, and was brought to The Marine Mammal Center October 22, 2011 for treatment. He showed no clinical signs of leptospirosis, but had an initial, moderately high anti-*Leptospira* antibody titer (log_2_ titer = 10). SUB2 was released back into the wild 3 weeks later but stranded again 3 months after the initial stranding event with flipper injuries and still no clinical signs of leptospirosis. He was never re-released and was adopted by the Navy July 11, 2012. Samples were provided for monitoring every 1-7 months for 44 months from Day 0, defined as the date of the first detected antibody titer, which was also the day of first stranding. It is possible that this animal experienced clinical disease prior to being monitored, however he never showed clinical signs consistent with leptospirosis while in rehabilitation or while at the U.S. Navy MMP. For both SUB1 and SUB2, the magnitude of the first detected anti-*Leptospira* antibody titers, and timing (autumn of 2011, during a major *Leptospira* outbreak in the wild sea lion population; Fig 2C&D and Fig 6A [23, 45]), suggest that exposure to *Leptospira* was recent – i.e. within weeks or months.

Samples collected from stranded animals (STRAND, CLINICAL and SUBCLINICAL) for this study were collected during routine clinical care at the rehabilitation centers and under their approved National Oceanic and Atmospheric Administration (NOAA), NMFS-Southwest Region Stranding Agreements under the authority of the Marine Mammal Protection Act. Samples collected from SUB1 and SUB2 at the U.S. Navy MMP were collected during their routine care and under U.S. Code, Title 10, USC 7524. The MMP houses and cares for a population of California sea lions in San Diego Bay (CA, USA). MMP is accredited by AAALAC International and adheres to the national standards of the U.S. Public Health Service Policy on the Humane Care and Use of Laboratory Animals and the Animal Welfare Act. During their clinical care, stranded animals received a variety of treatments which may have included, but were not limited to, subcutaneous fluids, antimicrobials, sedatives, and gastro-intestinal protectants.

### Sample Analysis

Serum agglutination testing (MAT) was performed at the California Animal Health and Food Safety (CAHFS) laboratory, Davis, California, or at the Centers for Disease Control and Prevention (CDC), Atlanta, Georgia, using live cultured *Leptospira* spp. (reference strains) to measure the serum anti-*Leptospira* antibody titers. Samples run at CAHFS were run against a 6 serovar panel and samples run at the CDC were run against a 2 or 19 serovar panel (as described in Prager et al. [28]). We only report MAT titer results against *L. interrogans* serovar Pomona as historically this is the strain that elicits the highest MAT titer in the majority of California sea lions tested [24] and it is the only serovar isolated from this species to date [27, 46]. Serum samples were tested at doubling dilutions starting from 1:100, and agglutination was read using dark field microscopy. Endpoint titers were reported as the highest dilution that agglutinated at least 50% of the cells for the strain tested [47]. Titer results were log transformed for ease of interpretation using the following formula: log_2_(titer/100) + 1, thus a titer of 1:100 = 1, 1:200 = 2, etc. Titers reported as <1:100 were set equal to 0 on both the log and regular scale. Throughout the paper “antibody titer” refers to this log transformed titer value. All animals with a detectable titer (i.e. ≥1:100) were considered seropositive and assumed to have been infected with *Leptospira* at some point.

Serum chemistry analyses of wild-caught sea lions and stranded sea lions from The Marine Mammal Center were performed on an ACE® Clinical Chemistry System (Alfa Wassermann, Inc., West Caldwell, New Jersey, USA), those of SUB1 were performed initially on either a VetTest® 8008 Chemistry Analyzer (IDEXX Laboratories, Inc., Westbrook, Maine, USA) or a Cobas 800 modular analyzer (Roche Diagnostics, Indianapolis, Indiana, USA). Once SUB1 and SUB2 were at the MMP, serum chemistry analyses were performed on a Roche Cobas 8000 system (Roche Molecular Systems, Pleasanton, CA, USA) by the Naval Medical Center in San Diego, CA.

We assessed leptospiral DNA shedding in urine using real-time polymerase chain reaction (PCR) as described in Wu et al., [48]. Because urine was collected under anesthesia, and anesthesia can pose a health risk to compromised animals, only apparently healthy wild-caught animals were caught and sampled, and of the STRAND animals, urine was collected only from those undergoing anesthesia for other reasons or during necropsy. Individuals shedding leptospiral DNA were considered infected and infectious as the primary mode of transmission of *Leptospira* is through either direct or indirect contact with leptospires shed in the urine of infected individuals [49].

### Data Analyses

#### Antibody Titer Kinetics

For data from each CLINICAL individual, we used linear regression to characterize how the log_2_-transformed antibody titer declined with time. We calculated the rate of antibody titer decline as the slope of the regression line and the t ½ as the negative inverse of the slope. Using these data we determined the median titer decay rate and t_1/2_ for the CLINICAL animals.

Visual inspection of the SUBCLINICAL data suggested a biphasic pattern of titer decline (Fig 2D) during the time that they were monitored, echoing findings from earlier work [29, 30]. Thus we used piecewise linear regression to estimate the titer decline rate and t_1/2_ of the first and second phases separately for SUB1 and SUB2. For each animal, we estimated the specific day that determined the change point of the regression by fitting models over a range of possible change points from 10 to 500 days and using the day that yielded the model with the minimum mean-squared error.

#### Shedding Duration

We used the median antibody titer decline rate (*r* = 0.058 log_2_ antibody titer units/day) of the CLINICAL animals (Table 2) as well as their median initial antibody titer (*t_i_* = 11) to calculate an approximate estimate of the lower bound of shedding duration (*D*) in days for each observed titer level *t* in PCR-positive sea lions using the following equation:

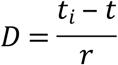

This is a lower bound because it ignores further shedding after the date of observation, and it neglects any shedding that occurred at titer levels > *t_i_*.

Similarly, using SUBCLINICAL animal biphasic decay rates, we calculated an approximate estimate of the duration of shedding (*D*_s_) if an animal started at an initial titer equivalent to the median initial antibody titer of the CLINICAL animals (*t_i_* = 11) and continued shedding until the animal became seronegative. Using the following equation we used the SUBCLINICAL specific decay rates (Table 2; *r*_1_ and *r*_2_) to estimate the durations of the initial and secondary phases for each animal, and the SUBCLINICAL specific titer at which the phase switch occurred (SUB1 *t_s_* = 2; SUB2 *t_s_* = 5) to mark the switch from initial to second phase decay rates:

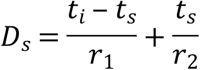

#### Renal Index

Blood urea nitrogen, creatinine, sodium, chloride and phosphorus are serum chemistry values known to change with leptospirosis-induced renal compromise [23]. We used principal components analysis (PCA) to derive a single measure of renal function from these five serum chemistry values, which we termed the renal index. PCA was performed in R using the command “prcomp” in the program “stats”[50]. BUN was log_10_ transformed and each individual serum chemistry measure was scaled to have unit variance prior to analysis. The dataset used included all longitudinal data (CLINICAL and SUBCLINICAL), as well as all WILD animals that were both seronegative for anti-*Leptospira* antibodies and negative for urinary leptospiral DNA shedding (i.e. the 0* group from Table 3). CLINICAL and SUBCLINICAL animals were included to capture the range of clinical compromise in infected animals from initial infection through recovery, and the subset of WILD animals was included to anchor the analysis with a group of apparently healthy, uninfected, unexposed animals. The first principal component (PC1) explained 54.8% of the variation in the data, and had factor loadings consistent with clinical reports of leptospirosis-induced disease (i.e. indicating elevated blood urea nitrogen, creatinine, sodium, chloride and phosphorus [23]). Therefore we used PC1 as the renal index to assess clinical severity of leptospirosis. Similar PCA results were found using just cross-sectional data (STRAND). To establish the range of values corresponding to healthy renal function, we computed the 95% interquantile range of renal index values (i.e. PC1) experienced by the apparently healthy WILD animals (−1.72 to 1.74). As values increased above this range, so did the degree of renal compromise.

Using the linear coefficients associated with each serum chemistry variable for PC1 from this analysis of longitudinal and WILD data, we calculated renal index values for all animals with serum chemistry results in the STRAND and WILD datasets.

#### Predicting Survival and Shedding

We used logistic regression to assess renal score as a predictor of survival in stranded animals at admission. We used the first sample available from all animals in the STRAND and CLINICAL groups with samples collected within 72 hours of admission (n=103) and that were categorized as leptospirosis cases based on clinical signs and serum chemistry values (BUN > 100 mg/dl, creatinine > 2mg/dl). Because we sought to assess the usefulness of this prediction as a tool for triaging animals in a rehabilitation center, antibody titer data were not included as only serum chemistry results would be available at this time.

In a separate analysis, we used multivariate logistic regression to assess predictors of leptospiral DNA shedding. Candidate predictors included serum anti-*Leptospira* antibody titer, renal index scores, and the interaction between these two variables. The dataset included all study animals for which we had PCR results, but only the first PCR result per individual. We used the “anova” command in the R package “stats” [50] to perform backward stepwise selection and the likelihood ratio method to include only variables that contributed significantly at the 0.05 level to the final model. Our final model included only antibody titer, so we then used the relationship between shedding and titer to predict the shedding status of the untested animals. To do this, we calculated the expected number of shedders amongst the untested animals at each observed titer level using the total number of untested animals and the probability of shedding at that titer level. We then randomly selected this expected number of animals from amongst the untested animals at that titer level and assigned them a positive shedding status. We performed logistic regression in R using the “glm” command in the package “stats” [50].

#### Comparing Renal Index Distributions

We used the Kolmogorov-Smirnov (KS) test to assess differences in distributions of renal index scores between groups of sea lions (CLINICAL, SUBCLINICAL, STRAND, and WILD) and within groups by anti-*Leptospira* antibody titer level. Because distributions were not continuous, we used the bootstrap Kolmogorov-Smirnov test “ks.boot” in the “Matching” package in R [51]. To achieve sufficient sample sizes, titers were collapsed into five groups, based on the titer kinetics observed in longitudinally followed animals (CLINICAL and SUBCLINICAL). The highest grouping included titers ≥ 11, consistent with the majority of the initial titers in CLINICAL animals (11/13 had titers ≥ 11). All CLINICAL animals were in the healthy renal index range by the time they had a titer of 8 and were released by the time their titer declined to 6, so these levels were used to define the ranges of the next two groupings: titers 9-10 to capture animals recovering from clinical disease, and 6-8 to capture the recovered, subclinical phase as seen in CLINICAL animals. Titer group 1-5 captured the longer-term subclinical phase, as seen in SUBCLINICAL animals. Titer group 0 captured seronegative animals.

#### 95% Confidence Intervals (CI)

95% CI in Fig 6 were calculated in R using binom.confint in the package “binom” using the Pearson-Klopper method [52]. 95% CI for Table 2 were calculated using normal approximations based on linear regressions for antibody titer kinetics.

#### Figures

All figures were made in R. Logistic regressions were plotted using logi.hist.plot in the package “popbio” [53], all other figures were made using ggplot in the package “ggplot2” [54].

### Ethics Statement

All California sea lion samples were collected under authority of Marine Mammal Protection Act Permits No. 932-1905-00/MA-009526 and No. 932-1489-10 issued by the National Marine Fisheries Service (NMFS), NMFS Permit Numbers 17115-03, 16087-03, and 13430. The sample collection protocol was approved by the Institutional Animal Care and Use Committees (IACUC) of The Marine Mammal Center (Sausalito, CA; protocol # 2008-3), the University of California Los Angeles (ARC # 2012-035-12), and the Marine Mammal Laboratory (Alaska Northwest 2013-1 and 2013-5). The Marine Mammal Center and the Marine Mammal Laboratory adhere to the national standards of the U.S. Public Health Service Policy on the Humane Care and Use of Laboratory Animals and the USDA Animal Welfare Act. UCLA and the U.S Navy Marine Mammal Program are accredited by AAALAC International and adhere to the national standards of the U.S. Public Health Service Policy on the Humane Care and Use of Laboratory Animals and the USDA Animal Welfare Act.

## Acknowledgements

We would like to thank the volunteers, veterinarians, biologists and staff from The Marine Mammal Center (Sausalito, CA), The Marine Mammal Care Center Los Angeles, the National Marine Mammal Laboratory Alaska Fisheries Service, Oregon and Washington Departments of Fish and Game, at the U.S. Navy Marine Mammal Program, the Año Nuevo State Park, and the University of California Santa Cruz’s Año Nuevo Reserve for their logistical support of this study and assistance with sample and data collection. In particular we would like to thank Carlos Rios, Lauren Palmer, Jeff Harris, Sharon Melin, Robert DeLong, Julia Burco, Patrick Robinson, Celeste Parry, Guy Oliver, and Pat Morris. We would also like to thank the Lloyd-Smith Laboratory at UCLA for useful discussions about the manuscript. This work was supported by the National Science Foundation Awards OCE-1335657 (https://www.nsf.gov/funding/pgm_list.jsp?org=OCE; JL-S, KCP) and DEB-1557022 (https://www.nsf.gov/funding/pgm_list.jsp?org=DEB; JL-S and KCP), the John H. Prescott Marine Mammal Rescue Assistance Grant Program (https://www.fisheries.noaa.gov/grant/john-h-prescott-marine-mammal-rescue-assistance-grant-program; JL-S), the Hellman Family Foundation (https://www.apo.ucla.edu/faculty-career-development/hellman-fellowship/hellman; JL-S), the US Department of Defense Strategic Environmental Research and Development Program Award RC-2635 (https://www.serdp-estcp.org/Funding-Opportunities/SERDP-Solicitations; JL-S, KCP), the De Logi Chair in Biological Sciences (https://www.apo.ucla.edu/academic-listings/endowed-chairs; JL-S), and the Research and Policy for Infectious Disease Dynamics (RAPIDD) program of the Science and Technology Directory, Department of Homeland Security, and Fogarty International Center, National Institutes of Health (https://www.fic.nih.gov/About/Staff/Pages/epidemiology-population.aspx JL-S, KCP, MGB). The funders had no role in study design, data collection and analysis, decision to publish, or preparation of the manuscript.

## Supporting Information Legends

**S1 Box.** Comparison of host-pathogen interactions based on the canonical ecological model of infectious disease dynamics in which individuals can be classified into four groups: susceptible (S), exposed (E), infected/infectious (I) and recovered (R), with more complex host-pathogen interactions.

**S2 Table.** Total number of sea lions with a given log_2_ antibody titer by year for wild-caught (WILD), stranded (STRAND) and subclinically infected (SUB1 and SUB2) sea lions. In parentheses are the number of animals shedding leptospires for each log_2_ antibody titer level over the total number of PCR tested animals.

**S3 Table.** Predicted shedding duration by titer level assuming a constant titer decay rate of 0.058 log2 antibody titer units/day (the median decay rate of the CLINICAL animals) and an initial titer of 11 (the median initial titer of the CLINICAL animals).

**S4 Data.** Raw data on used for analyses. Columns include animal ID, DataType (i.e. CLINICAL, SUBCLINICAL, STRAND, WILD), AdmitYear (i.e. the year an animal was caught – WILD – or admitted for rehabilitation - CLINICAL, SUBCLINICAL, STRAND), SampleYearMAT (year serum was collected for serum MAT), LogMAT (log_2_ MAT result), SampleYearChem (year serum was collected for serum chemistry analysis), RenalIndex (renal index score calculated as described in the manuscript), SampleYearPCR (year urine was collected for PCR, PCR (result of PCR analysis), SurvivalData (information whether the animal survived or died during rehabilitation; wild-caught animals were released after capture, therefore survival data was unknown – NA), DaySinceAdmission (the number of days between admission to rehabilitation and date of sample collection for analysis (MAT, PCR, serum chemistry), DaysSinceFirstMAT (the number of days since sample collection for the first MAT analysis).

